# Identification of genetic markers for the discrimination of *Bacillus thuringiensis* within the *Bacillus cereus* group, in the context of foodborne outbreaks

**DOI:** 10.1101/2022.04.22.489186

**Authors:** Arnaud Fichant, Arnaud Felten, Armel Gallet, Olivier Firmesse, Mathilde Bonis

**Affiliations:** Laboratory for Food Safety, University Paris-Est, French Agency for Food, Environmental and Occupational Health & Safety (ANSES), 94700 Maisons-Alfort, France; Université Côte d’Azur, CNRS, INRAE, ISA, France; Ploufragan-Plouzané-Niort Laboratory, viral genetics and biosafety Unit, French Agency for Food, Environmental and Occupational Health & Safety (ANSES), 22440 Ploufragan, France

**Keywords:** *Bacillus thuringiensis*, *Bacillus cereus*, foodborne outbreak, genome-wide association study

## Abstract

*Bacillus thuringiensis* (Bt), belonging to the *Bacillus cereus* (Bc) group, is commonly used as a biopesticide worldwide, due to its ability to produce insecticidal protein crystals during sporulation. The use of Bt, especially subspecies *aizawai* and *kurstaki*, to control pests such as Lepidoptera generally involves spraying mixtures containing spores and crystals on crops intended for human consumption. Recent studies have suggested that the consumption of commercial Bt strains may be responsible for foodborne outbreaks (FBOs). However, its genetic proximity to Bc strains has hindered the development of routine tests to discriminate Bt from other Bc, especially *Bacillus cereus sensu stricto* (Bc ss), also responsible for FBOs. Here, to develop tools for the detection and the discrimination of Bt in food, we carried out a genome-wide association study (GWAS) on 286 complete genomes of Bc group strains to identify and validate *in silico* new molecular markers specific to different Bt subtypes. The analyses led to the determination and the validation *in silico* of 128 molecular markers specific to Bt, its subspecies *aizawai, kurstaki* and four previously described proximity clusters associated with these subspecies. We developed a command line tool (https://github.com/afelten-Anses/Bt_typing) based on a 14-marker workflow for *in silico* Bt identification of a putative Bc genome with the aim of facilitating the discrimination of Bt from other Bc and between Bt subspecies, especially in the context of FBOs. Collectively, these data provide key elements for investigating Bc/Bt-associated FBOs and for monitoring Bt in food.

## 1. Introduction

*Bacillus cereus sensu lato*, or the *Bacillus cereus* group (Bc), is composed of Gram-positive, facultative anaerobic, spore-forming, ubiquitous bacteria. This group comprises at least nine species: *B. anthracis, B. cereus sensu stricto* (Bc s.s.), *B. thuringiensis* (Bt), *B. mycoides, B. pseudomycoides, B. weihenstephanensis, B. cytotoxicus, B. toyonensis* and *B. wiedmannii* (Miller et al., 2016). Furthermore, Bc members have very similar characteristics and highly conserved genomes (Ehling-Schulz et al., 2019). As a result, the taxonomic classification within the group is regularly revised, particularly due to the widespread use of whole-genome sequencing (WGS) which has led to higher resolution for the definition of new species (Carroll et al., 2021; Liu et al., 2017).

Bc possesses an extensive number of virulence genes and is well known to be an agent responsible for foodborne outbreaks (FBOs). According to the European Food Safety Authority (EFSA), in 2019, Bc represented the leading cause of FBOs due to bacterial toxins in Europe (EFSA, 2021). Bc can cause gastrointestinal disorders, such as diarrhea, vomiting, abdominal pain, or a combination thereof. The diarrhea is due to the ingestion of bacteria that produce enterotoxins, including three major pore-forming toxins: the non-hemolytic enterotoxin (Nhe) is found in approximately 85% to 100% of Bc strains, and hemolysin BL (Hbl) and cytotoxin K (CytK) are found in approximately 40 to 70% (Dietrich et al., 2021). In contrast, the cereulide toxin, produced in food by some vegetative Bc, can lead to emetic syndromes, which predominantly cause vomiting. The *ces* operon coding for the emetic toxin synthesis genes is located on a megaplasmid in a restricted range of Bc s.s. strains (Ceuppens et al., 2011). More rarely, Bc can also be responsible for extradigestive pathologies such as ocular and nosocomial infections (David et al., 1994; Glasset et al., 2018).

Unlike other Bc bacteria, during sporulation, Bt strains have the ability to produce parasporal crystals containing toxins, some of which are extremely toxic toward insect larvae; Bt strains are therefore widely used as biopesticides in organic and conventional agriculture (Roh et al., 2007). These crystals are composed of Cry and Cyt protoxins (also called δ-endotoxins) for which several hundred different haplotypes have been described so far (Crickmore et al., 1998). Their classification has been recently revised based on protein sequence identity (Crickmore et al., 2020). Upon ingestion of spores and crystals by larvae, larval digestive enzymes allow the release and activation of toxins from the crystals. Cry/Cyt toxins bind to specific receptors of the host midgut, forming pores in the cytoplasmic membrane of enterocytes, thereby triggering their death and creating breaches in the intestinal lining. In parallel, the favorable environment of the midgut supports the germination of spores and their entry in the internal milieu (Palma et al., 2014). Vegetative bacteria can proliferate, ultimately leading to septicemia and the death of the host. Four subspecies of Bt are commonly used in commercial products. Bt ssp. *aizawai* (Bta) and Bt ssp. *kurstaki* (Btk) and target Lepidoptera, Bt ssp. *morrisoni* (Btm) targets Coleoptera and Bt ssp. *israelensis* (Bti) is used against mosquitoes. Due to the ability of Bt to produce a large spectrum of insecticidal molecules, it is now considered as the leading microbial pesticide worldwide (Lacey et al., 2015).

Interestingly, Bc and Bt share some virulence genes, particularly those encoding enterotoxins (Dietrich et al., 2021). In addition, bacterial spores, in particular Bt spores, have the ability to persist in the environment and have been found on vegetables that have been treated with commercial products (Bonis et al., 2021; Frederiksen et al., 2006; Frentzel et al., 2020). Because Bc and Bt are genetically very closely related, there is no accurate test to distinguish between different Bc group members, meaning that discrimination between Bc s.s. and Bt is not straightforward. Nevertheless, optical microscopy can be used to detect parasporal insecticidal crystals (NF EN ISO 7932/Amd1), but this method requires expertise and several days to obtain analysis results. For instance, retrospective analyses in Canada (McIntyre et al., 2008) and France (Bonis et al., 2021) identified Bt in FBOs that were initially attributed to Bc. In a report dealing with the possible involvement of Bt in foodborne infection events, EFSA called for the development of a simple method to differentiate Bt from other Bc members (EFSA, 2016), and a fortiori the Bt strains used as pesticides. In addition, the issue of food monitoring and, in particular, the acceptable dose of Bc/Bt in food is regularly examined by health authorities. The development of new tools for the specific identification of Bt strains would facilitate and extend this monitoring, with the aim of preventing and limiting the emergence of new intoxication cases.

Genetic proximity between Bt and other members of the Bc group, especially Bc s.s. (Carroll et al., 2021; Ehling-Schulz et al., 2019), makes it difficult to develop identification methods; however, some genomic approaches may offer a solution. With the development of high-throughput sequencing (HTS), the rapid acquisition of complete genomes has been greatly facilitated. Based on pangenome analyses, recent studies have split the Bc group into three to five major clades (Bazinet, 2017; Ehling-Schulz et al., 2019), whereas a previous classification defined seven phylogenetic groups, named I to VII and associated with different levels of psychrotolerance (Guinebretière et al., 2008). According to the literature, the entire Bt species is spread across several of these clades or groups, but most Bt strains of interest with regard to food safety (e.g. isolated from pesticides or FBOs) show less divergence (Bonis et al., 2021) and cluster with other strains in phylogenetic group IV, clade 2. In an attempt to distinguish them, four proximity clusters (named a to d) have been defined using an SNP calling approach.

The genome-wide association study (GWAS) approach, originally developed for human studies, has been adapted for the analysis of microorganisms (Power et al., 2017). The GWAS approach can establish a statistical link between a feature polymorphism (named hereafter “trait”) and a molecular marker, such as a gene, a k-mer or a single nucleotide polymorphism (SNP). Several studies have already been carried out to correlate the presence of associated markers with a given phenotype, for example in the context of host-pathogen interactions (Vila Nova et al., 2019), persistence and resistance to antibiotics (Desjardins et al., 2016) as well as low-temperature growth phenotypes in food production (Fritsch et al., 2019). Nevertheless, to date, no studies have identified markers specific to bacterial species or subspecies for the investigation on FBO etiological agents.

In this study, we searched for specific marker sequences for the identification of Bt and some Bt groups of interest, among other Bc. We performed a pan-GWAS analysis on 230 complete genomes. A total of 249 markers strongly associated with various traits (Bt species, Bt subspecies and Bt clusters a to d) were identified. Among them, 128 showed potential informativeness as genomic markers after a TBLASTN validation step. For each designated trait, one marker or a combination of up to six markers could identify all the Bt in the studied dataset. We wrote a Python script using 14 markers to allow accurate and automated identification of Bt from a genomic assembly.

## 2. Materials and methods

### 2.1. Whole-genome sequencing

The genomic DNA of 78 Bt/Bc isolates was extracted and sequenced as described in Bonis *et al.* (2021). Briefly, the KingFisher Cell and Tissue DNA kit (ThermoFisher) was used to isolate genomic DNA. DNA purity and concentration were determined using a Nanodrop Spectrophometer and a Qubit fluorimeter, respectively. Global DNA integrity was visualized on a 0.8% agarose gel (Seakem GTG™ Agarose) after migration for 2 h at 90 V. Library preparation was carried out using the Nextera XT DNA Library Prep kit (Illumina) and 150 bp paired-end sequencing of isolated DNA was performed by the *Institut du Cerveau et de la Moëlle epinière*, using a Nextseq500 sequencing system (Illumina). An in-house workflow called ARtWORK v1.0 (Bonis et al., 2021; Mahamat Abdelrahim et al., 2019) was used to assemble reads with default parameters. The paired-end reads of the isolates are available in the PRJNA781790 BioProject on NCBI and the associated accession numbers are listed in Supplementary Table S1.

### 2.2. Dataset definition

A total of 286 genomes were used, corresponding to 215 FBO-associated Bc isolates, 18 commercial Bt strains (Bonis et al., 2021), and 53 Bc isolates of various origins, whose complete genomes were retrieved from the NCBI RefSeq database. In addition to the detection of parasporal insecticidal crystals under a microscope (NF EN ISO 7932/Amd1), Bt membership was confirmed by the presence of insecticidal protein-coding genes using the BtToxins_Digger pipeline v1.0.5 (Liu et al., 2021). Using R v4.0.1, a random draw was carried out to split the whole dataset into a study dataset (SD) and a validation dataset (VD) (Table S1), representing respectively 80% (*n* = 230) and 20% (*n* = 56) of the whole dataset. The SD was used as an input for the GWAS analysis, and the VD was used to validate the selected genetic markers. The SD included the genomes of 144 strains belonging to Bt species, especially from the subspecies *aizawai* (*n* = 56) and *kurstaki* (*n* = 57), and 86 genomes belonging to other Bc, including at least one assembly for each of the nine distinct representative species of the group (Table 1) and for each of the seven previously defined phylogenetic groups (Guinebretière et al., 2008) given in Table S1.

**Table 1.** Composition of the study dataset (SD).

| Species | Subspecies | No. of genome(s) | Representative genome | NCBI accession number |
| --- | --- | --- | --- | --- |
| *Bacillus anthracis* |  | 1 | Ames | NC_003997.3 |
| *B. cereus (sensu stricto)* (Bc) |  | 2 | ATCC 14579 | NC_004722.1 |
| *B. cytotoxicus* |  | 1 | NVH 391-98 | NC_009674.1 |
| *B. mycoides* |  | 1 | DSM 2048 | NZ_CM000742.1 |
| *B. pseudomycoides* |  | 1 | DSM 12442 | NZ_CM000745.1 |
| *B. thuringiensis* (Bt) | Bt ssp. *aizawai* | 56 | Leapi01 | AMXS00000000.2 |
|  | Bt ssp. *kurstaki* | 57 | HD-1 | NZ_CP004870.1 |
|  | Bt ssp. *israelensis* | 5 | AM6552 | NZ_CP013275.1 |
|  | Bt ssp. *morrisoni* | 1 | BGSC 4AA1 | NZ_CP010577.1 |
|  | other and/or unknown | 25 | ATCC 10792 | NZ_CP021061.1 |
| *B. toyonensis* |  | 1 | BCT-7112 | NC_022781.1 |
| *B. weihenstephanensis* |  | 1 | WSBC 10204 | NZ_CP009746.1 |
| *B. wiedmannii* |  | 1 | MM3 | NZ_CM000718.1 |
| other *B. cereus (sensu lato)* * |  | 77 |  |  |

\*genomes of Bc isolates collected from foodborne outbreaks, and for which, under sporulation conditions, no crystals were detected using optical microscopy.

### 2.3. Pangenome analysis

#### 2.3.1. Pangenome construction

An analysis was carried out to deduce the pangenome (i.e. core and accessory genes) from all 230 SD genomes (Fig. 1). The whole genome annotation of the assembly was performed using Prokka v1.13.3 (Seemann, 2014) with default parameters and GFF3 files were provided to Panaroo v1.2.3 (Tonkin-Hill et al., 2020) as input. Panaroo was run using the strict mode with default identity parameters and length difference thresholds (98%) for initial clustering of protein sequences with CD-HIT v4.8.1 (Li and Godzik, 2006). Then, close clusters were collapsed into putative families when they had at least 70% sequence identity. The assignment of a gene cluster to a core genome was defined by the presence of the gene in at least 99% of the genomes (*n* = 228). As output, Panaroo produced a presence/absence matrix, used for the pan-GWAS analysis.

**Fig. 1.**
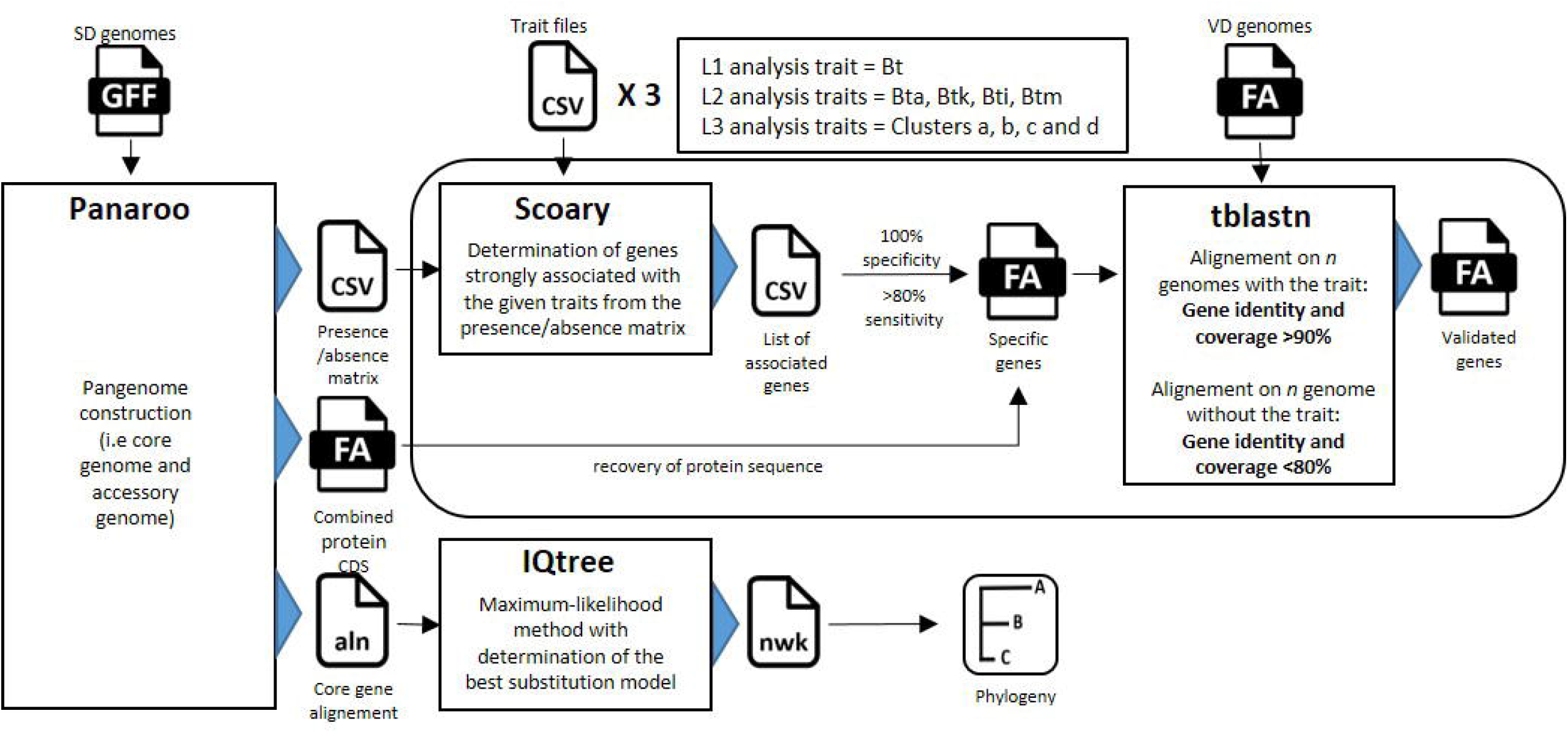
Flowchart of the pan-GWAS analysis. Pangenome construction was first performed using Panaroo (Tonkin-Hill et al., 2020) to obtain an estimation of the core and accessory genomes of the study dataset (SD). Then, the presence/absence matrix output from Panaroo was used for the GWAS analysis performed using Scoary, thereby providing a list of associated genes for each trait. Only genes with at least 80% sensitivity and 100% specificity for the SD were selected. A TBLASTN validation step of the selected genes was performed after recovering the protein sequences from Panaroo. The genes were validated when the respective sequence criteria of coverage and identity were met and when exhibiting at least 80% sensitivity and 100% specificity for the validation dataset (VD). The core-gene alignment obtained from Panaroo was used to construct the phylogeny of the 230 genomes, using IQtree (Nguyen et al., 2015).

#### 2.3.2. Genotype association

A three-level pan-GWAS analysis was performed to associate the presence or absence of a gene with a particular phenotype or trait. The first GWAS analysis (named L1 for Level 1) was conducted to search for genes specific to Bt species and the second analysis (named L2 for Level 2) for the Bt subspecies: Bt ssp. *aizawai* (Bta), Bt ssp. *kurstaki* (Btk), Bt ssp. *israelensis* (Bti) and Bt ssp. *morrisoni* (Btm). Then, specific genes for the SNP proximity clusters a, b, c and d within Bt subspecies (a, b for Bta and c, d for Btk) were investigated (analysis L3 for Level 3). These clusters were identified in a previous SNP study (see Bonis *et al.* 2021) and involve Bta or Btk strains that show strong genetic proximity with commercial strains used as insecticides. After targeting traits for each GWAS analysis, a gene search was carried out within the SD. Scoary v1.6.16 (Brynildsrud et al., 2016) was run to identify statistically robust genes (Fisher’s test) associated with each of the designated traits. For each gene, 1000 replicate permutations were conducted and *p*-values were adjusted by applying the Bonferroni correction (Abdi, 2007) and the Benjamini-Hochberg procedure (Wright, 1992). Only genes with *p*-values < 0.05, 100% specificity and at least 80% sensitivity for the screened traits were selected for the validation steps. In case of cluster-specific gene investigation, GWAS analysis (L3) was performed only on SD genomes with clear cluster attribution (*n* = 104), due to the impossibility to determine trait-specific genes when performed against all SD genomes. For each selected gene cluster, a protein sequence was extracted and tested as a candidate marker.

#### 2.3.3. Core genome phylogeny

Core-genome sequences from Panaroo were aligned using MAFFT v7.471 (Katoh et al., 2002). A phylogenetic tree of the SD was built with the maximum-likelihood method using iQtree v1.6.9 (Nguyen et al., 2015) and the best substitution model GTR+F+R10 was determined with ModelFinder (Kalyaanamoorthy et al., 2017) on 286 DNA models. The core genome phylogeny was visualized using Phandango v1.3.0 (Hadfield et al., 2018), alongside pangenome metadata. Selected marker presence/absence information within the SD was added to the phylogenetic tree using iTol v6.1.2 (Letunic and Bork, 2019).

#### 2.3.4. *In silico* validation of genetic markers

To certify the reliability of the selected markers, we carried out a TBLASTN v2.7.1+ against the VD to validate candidate marker sequences meeting the required conditions. For each cluster associated with a given marker, a protein sequence was retrieved from the Panaroo output and screened for in the VD genomes. Markers were selected only when sequence coverage and identity were strictly higher than 90% compared with genomes displaying the same trait (intragroup), and when the sequence coverage and identity were strictly lower than 80% compared with genomes not associated with the specific trait (intergroup). This extra step made it possible to test the pan-GWAS selected markers *in silico* against a new dataset of genomes, and to confirm the sensitivity and the specificity of the markers. To enhance the identification of a genotype, a combination of validated makers associated with the best sensitivity was defined. Predicted genomic localization (i.e. chromosome vs. plasmid) of the gene encoding for the protein selected as a marker was carried out with TBLASTN using protein sequences and a Bt plasmid database (*n* = 342) collected from PLSBD (Galata et al., 2019).

## 3. Results

### 3.1. *Bacillus cereus* group pangenome

To identify specific Bt markers associated with different traits of interest using a pan-GWAS approach with Scoary, the pangenome of the Bc group was inferred using Panaroo on SD, resulting in a total of 39,021 clusters of genes. The core genome of the 230 Bc complete genomes was composed of 1854 genes (representing about 5% of all genes), leaving 37,167 accessory genes (Fig. 2). Among the accessory genes, 1695 were considered soft core genes (presence in 95 to 99% of all genomes), 4358 shell genes (presence in 15 to 95% of all genomes) and 31,114 cloud genes (presence in less than 15% of all the genomes). A rarefaction curve of the 230 genomes (Supplementary Fig. S1) suggests the existence of an open pangenome, as previously assumed for Bc (Bazinet, 2017; Zwick et al., 2012). The presence-absence matrix showed that the *B. cytotoxicus* population, which roots the tree, seemed to possess a reduced number of accessory genes. In contrast, the Bta and Btk subspecies, whose genomes represent highly conserved populations, displayed a greater number of accessory genes.

**Fig. 2.**
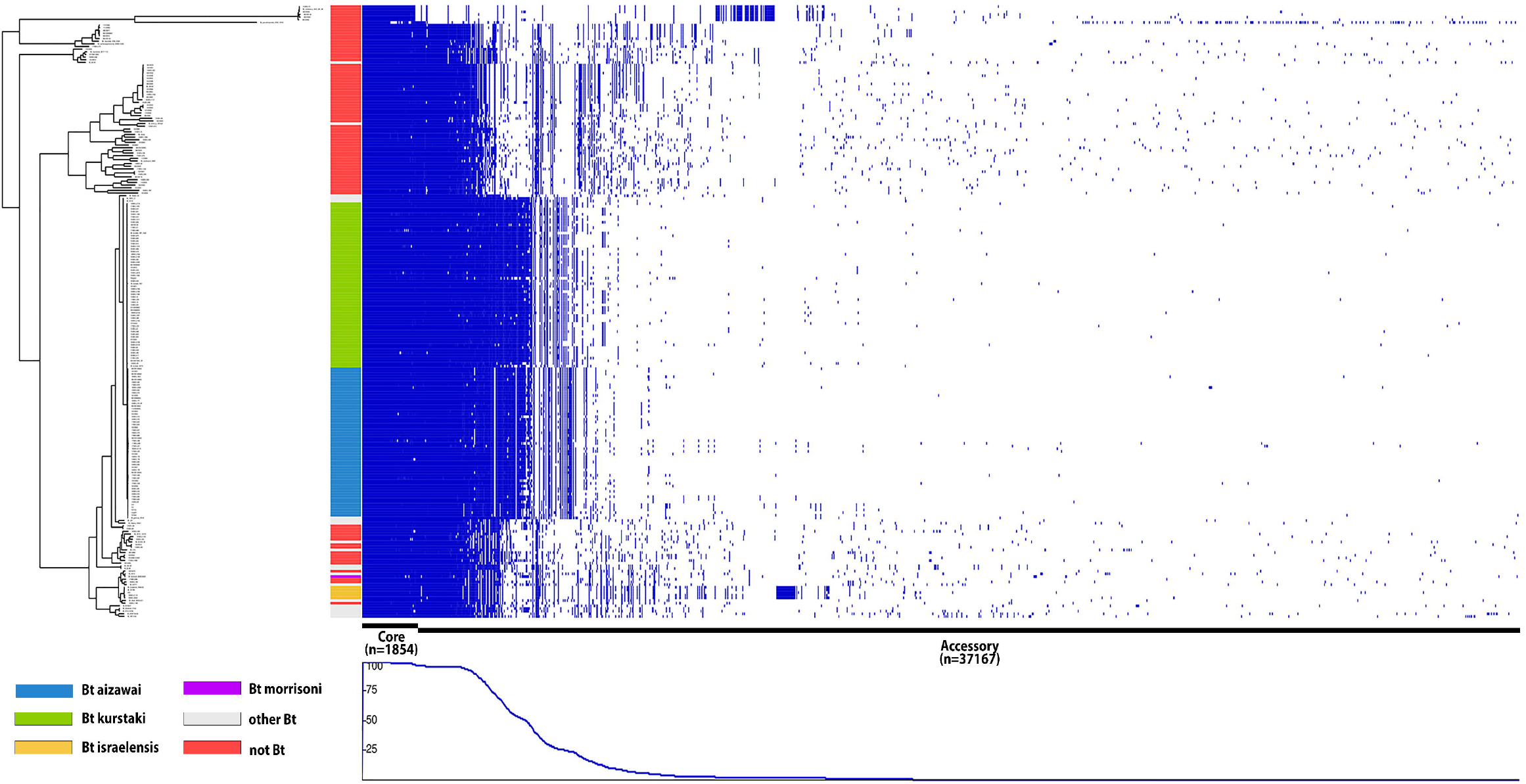
Phylogenetic tree of 230 *Bacillus cereus* (Bc) genomes alongside the gene presence/absence variation matrix. The maximum-likelihood phylogeny (GTR+F+R10 substitution model) was performed with IQtree (Nguyen et al., 2015) using the Panaroo (Tonkin-Hill et al., 2020) core-gene alignment of the study dataset (SD) genomes and visualized with Phandango (Hadfield et al., 2018). The presence-absence matrix obtained with Panaroo identified 1854 core genes (present in 99% of SD genomes) and 37,167 accessory genes for the 230 Bc genomes.

### 3.2. Pan-GWAS analysis

#### 3.2.1. Gene-based GWAS

We used Scoary to associate specific genes with the presence or absence of specific traits with a GWAS analysis. Three GWAS analyses were run to identify a large number of candidate genes for each trait (Table 2). A total of 1304 genes were significantly associated with the traits of interest. Interestingly, the majority of the identified genes were associated with subspecies *israelensis* and *morrisoni* in the GWAS L2 analysis, with 809 and 246 genes, respectively. However, this large number of associated genes can be explained by the low number of genomes available for these two subspecies in the dataset reducing the analytical strength. Given the low diversity in the dataset for these subspecies, we did not pursue the search for specific markers for these subspecies.

**Table 2.**
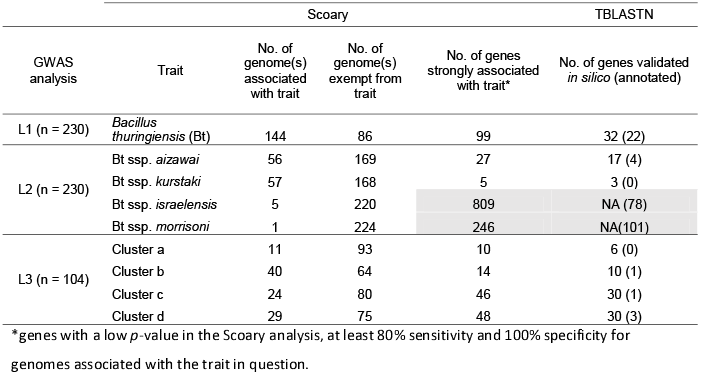
Summary results of the validation of specific markers.

Among the other 249 candidate genes, 131 genes met the specificity and sensitivity requirements for the L1 and L2 GWAS analyses, corresponding respectively to the Bt species trait and the Bt subspecies traits of the two remaining subspecies (i.e. Btk and Bta). Accordingly, 99 genes were specific to the Bt species and 27 to the Bta subspecies. Interestingly, only 5 genes were specific to Btk suggesting relatively high diversity within the Btk subspecies. For the L3 GWAS analysis performed on a smaller panel of genomes, 118 genes were identified among the four traits corresponding to clusters a, b, c and d with 10, 14, 46 and 48 genes, respectively.

#### 3.2.2. Validation of GWAS results

We then validated the markers selected using GWAS by comparing them *in silico* to the VD using TBLASTN. For each selected candidate gene associated with a specific trait in the GWAS analysis, a protein sequence was extracted and compared to all VD genomes that shared or did not share the trait in question. Out of the 249 highly associated genes obtained with Scoary, 128 protein sequences met the criteria of coverage, specificity and sensitivity of the TBLASTN analysis (Table 2). All validated marker sequences had a sensitivity range between 80% and 100% on both SD and VD datasets. Then, when a single marker was not sufficient, combinations of markers were tested to identify all the associated genomes for each trait. Table 3 lists 14 markers associated with low *p*-values and whose combination could detect all genomes of a given trait (i.e. 100% sensitivity).

**Table 3.** List of 14 genes with strong evidence and best combinations for *Bacillus thuringiensis* (Bt) trait identification.

| GWAS analysis | Gene name | Replicon | Annotation | UniprotKB | Trait | Marker Sensitivity (%) | Cumulative Sensitivity (%) | p-value | NCBI accession number |
| --- | --- | --- | --- | --- | --- | --- | --- | --- | --- |
| L1 | *cwlA* | Chromosome | N-acetylmuramoyl-L-alanine amidase CwlA | P24808 | Bt | 93.8 | **93.8** | 1.72E-51 | WP_021728236.1 |
|  | *intQ* | Chromosome | Putative defective protein IntQ | P76168 | Bt | 91.7 | **94.4** | 1.64E-48 | WP_000237488.1 |
|  | group_3916 | Chromosome | Hypothetical protein | N/A | Bt | 88.9 | **95.1** | 3.29E-47 | WP_000858032.1 |
|  | group_20749 | Chromosome | Hypothetical protein | N/A | Bt | 86.8 | **97.2** | 6.36E-45 | WP_042596929.1 |
|  | group_20361 | Plasmid | Hypothetical protein | N/A | Bt | 84.7 | **98.6** | 8.43E-43 | WP_002101540.1 |
|  | *sdpR* | Plasmid | Transcriptional repressor SdpR | O32242 | Bt | 84.0 | **100.0** | 4.00E-42 | WP_000998670.1 |
| L2 | *apr* | Chromosome | Subtilisin | P04189 | Bta | 100.0 | **N/A** | 9.10E-53 | WP_021728520.1 |
|  | group_27293 | Plasmid | Hypothetical protein | N/A | Btk | 98.2 | **98.2** | 5.83E-52 | WP_003273526.1 |
|  | group_27336 | Plasmid | Hypothetical protein | N/A | Btk | 89.5 | **100.0** | 1.05E-43 | WP_001293418.1 |
| L3 | group_10114 | Chromosome | Hypothetical protein | N/A | Cluster a | 100.0 | **N/A** | 4.48E-15 | WP_000415284.1 |
|  | *rapF* | Chromosome | Response regulator aspartate phosphatase F | P71002 | Cluster b | 92.5 | **92.5** | 4.82E-25 | WP_050062578.1 |
|  | group_20667 | Plasmid | Hypothetical protein | N/A | Cluster b | 82.5 | **100.0** | 1.34E-20 | WP_131256056.1 |
|  | *lexA* | Chromosome | LexA repressor | P31080 | Cluster c | 100.0 | **N/A** | 4.31E-24 | AHZ54004.1 |
|  | *clpP1* | Chromosome | ATP-dependent Clp subunit 1 | B0B803 | Cluster d | 100.0 | **N/A** | 2.13E-26 | WP_000791073.1 |

#### 3.2.3. Functions and distribution of specific markers

To predict whether the genes coding for the markers were localized or not on a plasmid, a TBLASTN alignment on a Bt plasmid database was carried out (Table 3). When the function was known, the closest protein identified in Uniprot (The UniProt, 2017) was indicated. The Bt species can thus be identified with a combination of only six markers. For example, the *cwlA* gene located in an operon along with *cwlB*, encodes a peptidoglycan hydrolase, known for its role in cell lysis during Bt spore release (Yang et al., 2013). Regarding the markers that emerged from the GWAS L2 analysis for Bt subspecies, all Bta genomes (*n* = 56) could be identified based on the *apr* gene, which encodes a specific chromosomal alkaline serine protease, also called subtilisin in *Bacillus subtilis* (Park et al., 1989). For Btk, two genes encoding hypothetical proteins (group_27293 and group_27336) were needed to identify the 57 genomes present in the SD. Moreover, both genes were located on plasmids suggesting the presence of a common plasmid in these subspecies. Regarding the cluster-specific markers identified with the GWAS L3 analysis, clusters a, c and d could each be identified with a single chromosomal marker, respectively group_10114 encoding a hypothetical protein, the repressor *lexA*, involved in the SOS system (Au et al., 2005) and the gene encoding a subunit of Clp protease (Krüger et al., 2000). Cluster b, like Btk, needed two markers for the identification of all genomes: the *rap*F gene, involved in bacterial competence (Bongiorni et al., 2005), combined with a plasmid gene of unknown function (group_20667). For all hypothetical protein-coding genes, we attempted to annotate them with the 3D structure using SWISS-MODEL (Waterhouse et al., 2018). However, it was not possible to assign new putative functions to them (Supplementary Table S3).

Phylogenetic reconstruction of the 1854 core genes from the 230 genomes (Fig. 3) showed a clear distinction between most Bc genomes, including Bt genomes, and five genomes of the *B. cytotoxicus* species, which is known to be the most distant group member (Ehling-Schulz et al., 2019). In contrast, the genomes of the two subspecies Btk and Bta were located on the outermost branch of the tree compared with *B. cytotoxicus* genomes. Unsurprisingly, other Bt subspecies or unknown subspecies genomes were scattered around the tree. Some of them, such as MTC28 (NCBI accession: NC_018693.1) and Bt *finitimus* YBT020 (NCBI accession: NC_017200.1) were phylogenetically very distant.

**Fig. 3.**
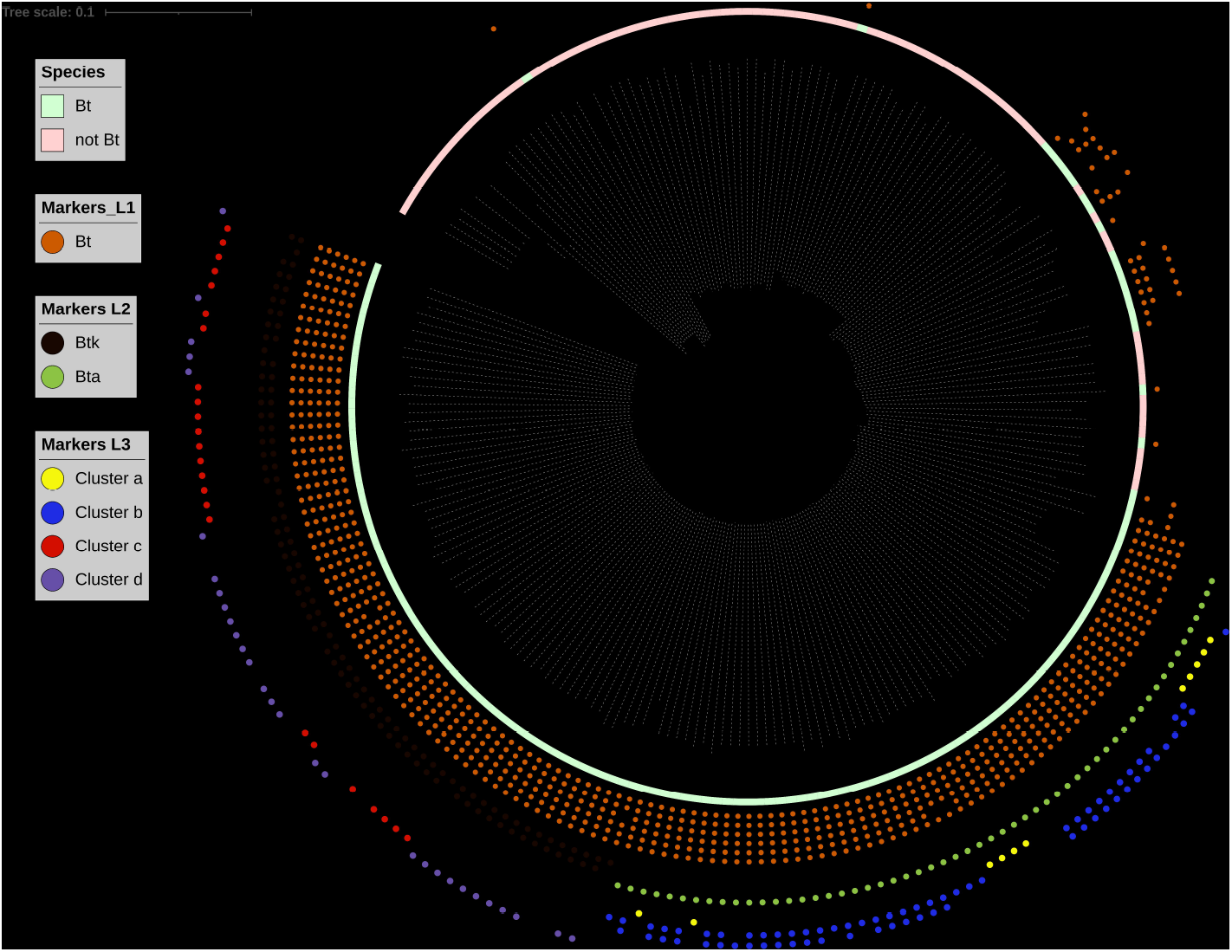
Core genome phylogeny of 230 *Bacillus cereus* (Bc) genomes. The maximum-likelihood phylogeny (GTR+F+R10 substitution model) was performed with IQtree (Nguyen et al., 2015) using a Panaroo (Tonkin-Hill et al., 2020) core-gene alignment of the study dataset (SD) genomes. Visualization and annotation were performed using iTol (Letunic and Bork, 2019). Filled circles indicate the presence of the 14 Bt markers for the successive GWAS analysis levels (L1, L2 and L3) in the corresponding SD genomes.

The 14 Bt-specific markers identified with the three levels of GWAS analysis and validated by TBLASTN were mapped on the phylogenetic tree (Fig. 3). Despite the great distance between some Bt strains, a combination of six markers can identify all of them. Similarly, with two markers specific to Btk and cluster b, all associated genomes can be retrieved. Only one marker was needed for the identification of Bta, clusters a, c and d.

#### 3.2.4. Workflow for Bt identification

Based on the 14 selected markers, a workflow was developed to distinguish Bt from the genome of a putative Bc isolate (Fig. 4). We wrote an in-house script in Python 3 to automate the screening of the markers in genome assemblies. Using BLASTX, the script detects the markers for Bt, Bt subspecies and Bt clusters using the same thresholds as the TBLASTN validation (90% identity and coverage) in the user-input genomes. The output indicates the presence (+) or absence (-) of all trait-associated markers for each genome query. Then, according to the workflow pathway, based on the presence/absence of the markers within the genome, a putative Bt identification is given. The script is freely available at https://github.com/afelten-Anses/Bt_typing.

**Fig. 4.**
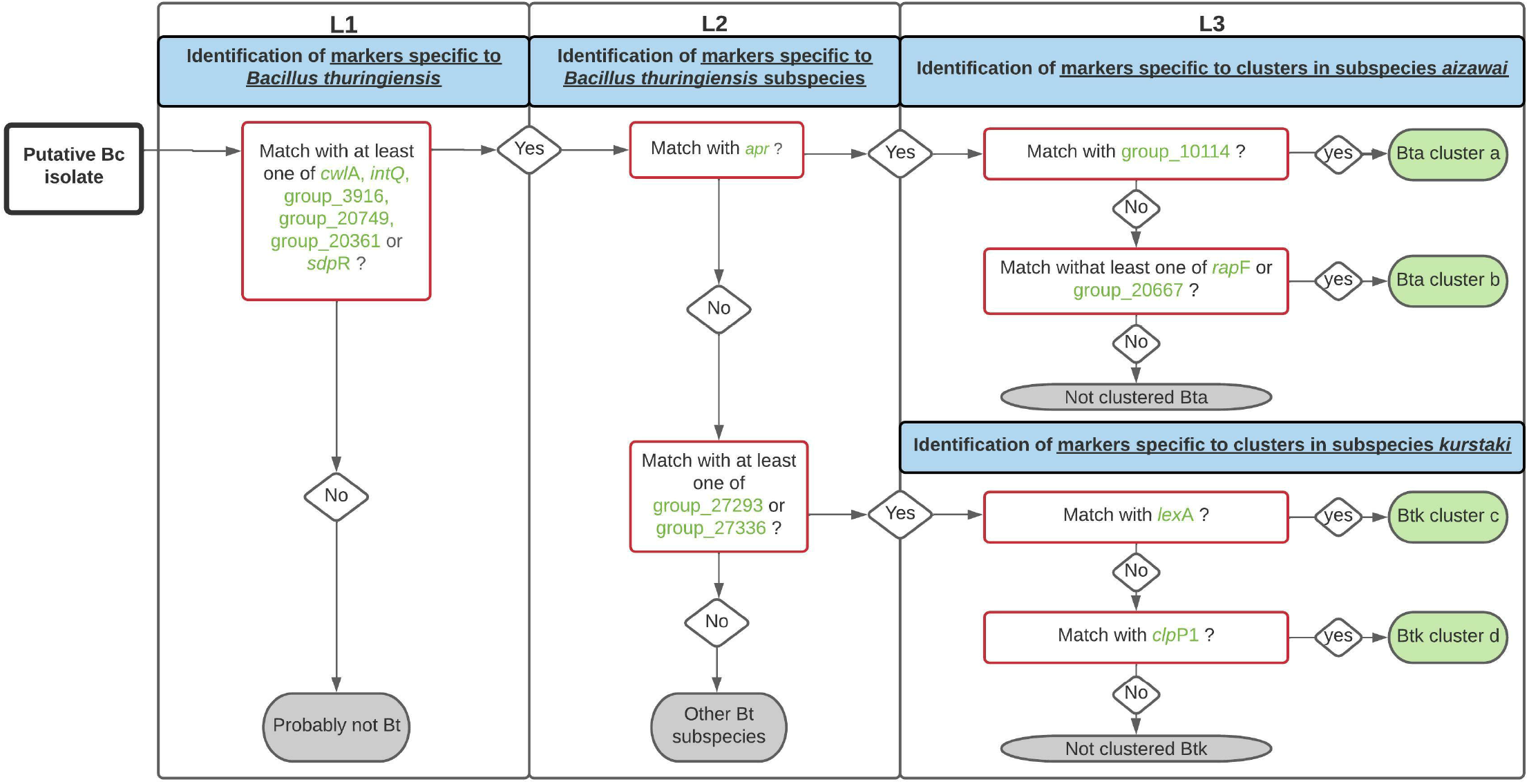
Workflow for *Bacillus thuringiensis* (Bt) identification. The proposed decision tree is used to determine if a putative *Bacillus cereus* (Bc) isolate belongs to the Bt species (L1), to the Bt ssp. *kurstaki* (Btk) or *aizawai* (Bta) subspecies (L2) and to the clusters a, b, c, and d (L3), based on the 14 marker protein sequences previously identified (see Table 3). A marker is considered present when its sequence coverage and its identity are both higher than 90% in the assembly tested. The workflow was automated using a Python script available online (https://github.com/afelten-Anses/Bt_typing).

## 4. Discussion

To differentiate Bt from other members of the Bc group and especially in the context of food poisoning, we analyzed a large database of Bc to identify genetic markers specific to Bt. The construction of the Bc pangenome from 230 Bc genomes showed that it was composed of 39,021 protein-coding genes and that the core genome corresponded to 1854 genes. These results differ somewhat from those previously published for the Bc pangenome. A previous pangenome study of 114 Bc highlighted 59,989 genes, but only 600 were described as core genes (Bazinet, 2017). Normally, increasing the number of genomes in a dataset can lead to a decrease in the core genome size along with an increase in the accessory genome size. However, the difference of tools used for pangenome deduction here can explain this large difference in gene numbers, as already highlighted in a genome-wide analysis of *Clostridium difficile* pangenome (Knight et al., 2021). For instance, the previous popular pangenome analysis software, named Roary (Page et al., 2015), uses a default threshold of 95% identity (versus 98% with Panaroo) to create clustered protein sequences. Nonetheless, Panaroo, unlike Roary, clusters potential close family genes together. Moreover, Roary does not take assembly and annotation errors into account, thereby resulting in an overestimation of the total gene number and thus a reduction of the core genome size, especially for datasets with high diversity as in the Bc group.

In addition, many Bt genomes, especially from the Btk and Bta subspecies, used for this study comprised a large number of accessory genes, which can be explained by the high diversity of the genomes in these subspecies in the dataset and/or the presence of mobile genetic elements, particularly plasmids. Several studies have already highlighted that the number of plasmids is high in Bt in comparison to Bc (Fayad et al., 2019). A comparative genomic analysis study has shown that Bt strains with high insecticidal potency harbor genes promoting infection, immune evasion and nutrient access that may play a key role in entomopathogenicity and host adaptation (Liu et al., 2015). Toxin-carrying plasmids, which represent an important part of the Bt plasmid pool, have also been shown to be involved in cellular functions such as germination, sporulation and horizontal gene transfer (Fayad et al., 2019; Gillis et al., 2018). For the Bti and Btm subspecies, for which only a limited number of genomes were included in this study, no gene validation step could be performed to define specific markers. Nevertheless, our analysis led to the identification of genes only present in a restricted population of Bti (18SBCL211A, 18SBCL484A, Bt_israelensis_4Q1, Bt_israelensis_HD789, Bt_israelensis_AM6552) or in one strain of Btm (Bt_morrisoni_BGSC 4AA1), whose specificity deserves further investigation.

A total of 249 candidate genes were selected based on their sensibility and specificity for a given trait. As recommended for GWAS analyses (San et al., 2019), the results obtained were validated by testing them on a test dataset (here, the VD). The fact that almost 50% of the selected markers did not pass this step demonstrates the importance of testing results. To extend identification to other traits or Bt genomes, marker selection preferentially targeted genes with a chromosomal location and a known or predicted function. Of the 128 validated genes, only 21 had predicted functions, annotated with Prokka (Seemann, 2014) (Supplementary Table S3). At least one gene associated with a known function and located on the chromosome was identified for L1- and L2-associated traits (Bt species and subspecies, respectively), with the exception of Btk. However, we showed that Btk could be identified based on the combination of two plasmid markers, which underlines the importance of plasmid content, especially for the discrimination of closely related Bt strains.

The phylogenic visualization of the SD dataset illustrates the challenge — even when using genomic approaches — of differentiating Bt populations from each other or from other Bc group members, due to their close genetic proximity (Bazinet, 2017; Carroll et al., 2021). For example, L3 analysis performed for cluster identification could not be performed on the whole SD. The impossibility of identifying specific genes revealed a limitation of gene-based GWAS when comparing extremely close groups. Fortunately, the sequential use of a combination of markers allows the use of cluster-specific markers after confirmation of Bt membership and then one of two subspecies, Bta or Btk. Furthermore, the use of a highly diverse dataset in this study allowed the identification of specific and high sensitivity markers. For example, among the Bt markers identified, the *cwlA* gene was found in 93.8% of SD Bt genomes (*n* = 144), making it a very reliable marker compared with the previous proposed detection system (Chang et al., 2003; Chelliah et al., 2019; Dzieciol et al., 2013). Furthermore, this previous system did not include commercial Bt strains, unlike the analyses performed for this study. Nevertheless, it is noteworthy that the identified markers refer to a specific dataset, meaning they may not allow the identification of the entire Bt species. In addition, we cannot exclude the existence of some non-sequenced strains, in particular divergent strains not included in our dataset that may not possess one or more of the selected markers. However, the workflow developed here can easily be adapted with the addition of new markers or modification of existing markers, to detect additional Bt strains of interest.

The 2016 EFSA report (EFSA, 2016) highlighted the need for the development of new methods to differentiate Bt from Bc s.s. Here, to differentiate Bt from other members of the Bc group, particularly in the context of FBOs, our work shows that a combination of six markers can identify Bt species, three for two major subspecies of interest in food safety, and five for the proximity clusters of pesticide strains frequently used in agriculture. Based on these 14 markers, we developed a new tool, associated with a workflow (Fig. 4) and a script, to predict the identity putative Bt among Bc isolates. Moreover, the gene-based GWAS approach developed here demonstrated that the four proximity clusters associated with commercial Bta or Btk strains likely belong to Bt (Bonis et al., 2021). Currently, the workflow cannot be used to distinguish certain Bt strains (particularly the insecticide strains belonging to the *kurstaki* subspecies) within the same cluster. Gene-based approaches are limited in their specific identification of commercial Bt strains used in agriculture. New approaches, notably based on SNP calling, can be conducted to search for differentiation methods at the strain level. In this perspective, our workflow can be easily updated and optimized, if necessary.

With the routine use of sequencing methods in laboratories and the significant development of high-throughput sequencing techniques, the use of computational tools as a complementary method for the identification of bacterial species may prove to be a valuable asset, especially in the context of food poisoning. This complementary identification method can be used to quickly assign a Bc strain to the Bt species, as well as to challenge false positives and false negatives resulting from microscope searches for protein crystals, possibly due to ambiguous phenotypes, misinterpretation of results, or loss of plasmids carrying the *cry* genes. With different levels of analysis, possible assumptions on the origin of the isolates can be made. qPCR detection systems hold promise for the specific detection and quantification of Bt in food matrices, an important, as-yet-unattained goal for monitoring Bt in food.

## Acknowledgements

This project was funded and supported by ANSES (PPV project “OSABt”) via the tax on sales of plant protection products. The proceeds of this tax are allocated to ANSES to finance the development of a system to monitor the adverse effects of plant protection products, called “phytopharmacovigilance” (PPV), established by the French Act on the Future of Agriculture, Food and Forestry of 13 October 2014. A. Fichant was funded by the Consortium Biocontrol (BACILLUS project).

We thank all the district veterinary and food analysis laboratories for carrying out Bc detection and transmitting isolates to the Food Safety Laboratory.

## Supplementary data

**Supplementary Fig. S1**. Rarefaction curve of gene diversity as a function of the number of samples generated with non-parametric incidence-based estimator jack1 (Burnham and Overton, 1978).

**Supplementary Table S1**. List of the 286 genomes used in this study. The genome set was divided into two datasets, a study dataset (SD) (*n*=230) and a validation dataset (VD) (*n*=56). ^a^ Clusters were determined by SNP calling, using iVARCall2 (Bonis et al., 2021). ^b^ All the sequencing data used in this study are associated with two BioProjects: PRJNA547495 (Bonis et al., 2021) and PRJNA781790 (this study). The attribution to panC groups was performed after partial sequencing of *panC* gene, and according to Guinebretière *et al.*, (Guinebretière et al., 2008). Abbreviations: FBO=foodborne outbreak.

**Supplementary Table S2.** Search for 3D structure prediction for workflow markers of unknown function. The search was performed using SWISS-PROT. ^a^GMQE (global model quality estimation) is a quality estimation that combines properties from the target–template alignment and the template structure. ^b^QMEAN is a composite estimator based on different geometrical properties and provides both global (i.e. for the entire structure) and local (i.e. per residue) absolute quality estimates on the basis of one single model. ^c^QSQE (quaternary structure quality estimate) score is a number between 0 and 1, reflecting the expected accuracy of the interchain contacts for a model built based on given alignment and template.

**Supplementary Table S3**. List of the 21 annotated genes found with GWAS analysis and validated using TBLASTN.

