## Supplementary figures and images for "Identification of genetic markers for the discrimination of *Bacillus thuringiensis* within the *Bacillus cereus* group, in the context of foodborne outbreaks"

### Supplementary Fig S1

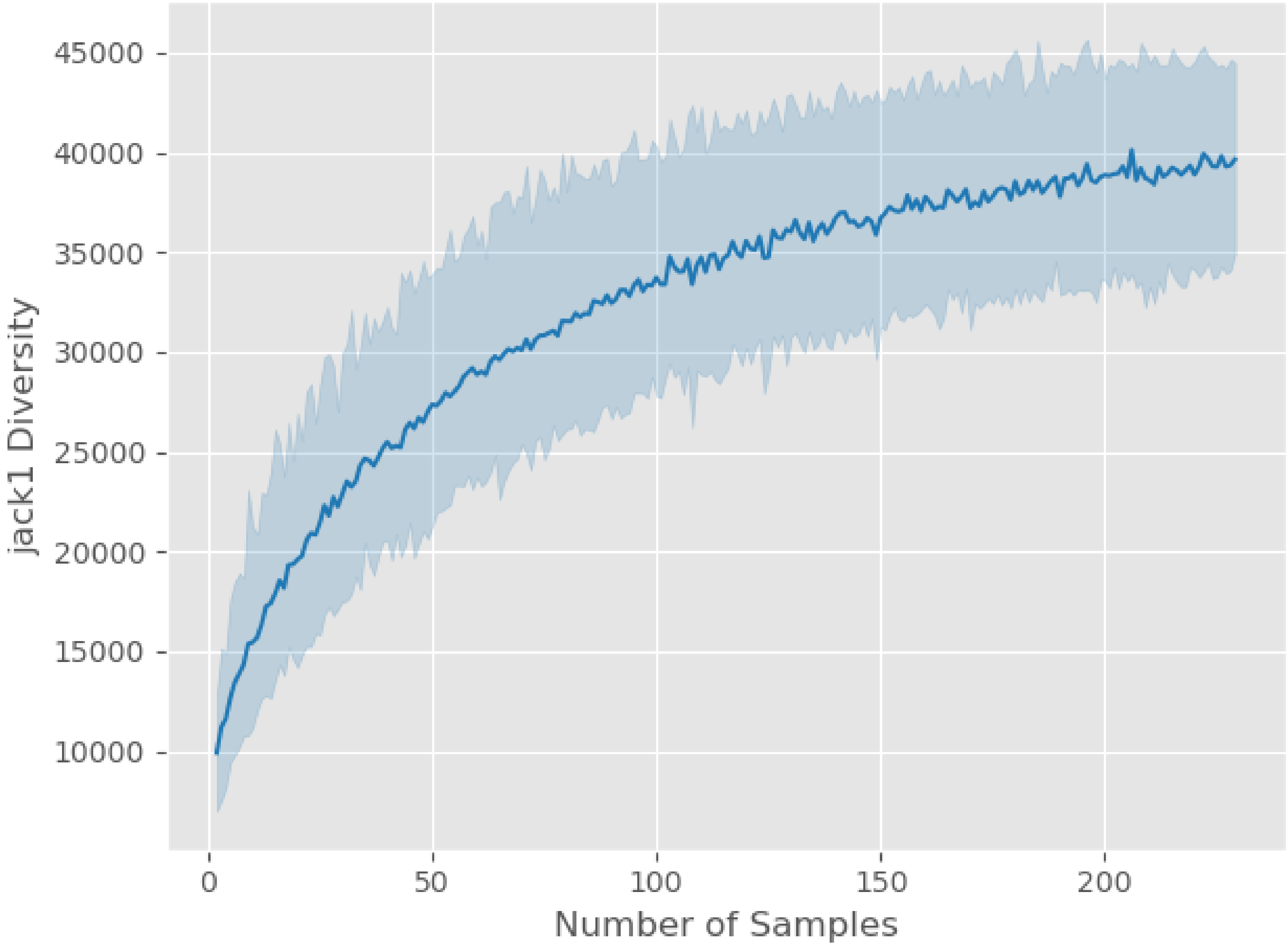
